# Patient-derived lymphomoids preserve the tumor architecture and allow to assess response to therapies in lymphoma

**DOI:** 10.1101/2024.04.15.589480

**Authors:** Albert Santamaria-Martínez, Justine Epiney, Divyanshu Srivastava, Daniele Tavernari, Marco Varrone, Dina Milowich, Igor Letovanec, Thorsten Krueger, Rafael Duran, Giovanni Ciriello, Anne Cairoli, Elisa Oricchio

## Abstract

The efficacy of anti-cancer therapies depends on the genomic composition of the tumor, its microenvironment, spatial organization, and intra-tumor heterogeneity. B cell lymphomas are a heterogeneous group of tumors emerging from B cells at different stages of differentiation and exhibiting tumor-specific interactions with the tumor microenvironment. Thus, to measure response to therapy in lymphoma, it is critical to preserve the tumor composition and functional interactions among immune cells. Here, we developed a platform to maintain small fragments of human lymphoma tissue in culture for several days and use them to test response to therapies. We collected 25 patient samples representative of different lymphoma subtypes and established *ex vivo* tissue fragments that retained histological, cellular, and molecular characteristics of the original tissue, here referred to as *lymphomoids*. Using lymphomoids, we tested sensitivity to several clinically approved small molecule inhibitors in parallel and examined tissue remodeling upon treatment. Importantly, when this information was available, we showed that sensitivity to therapy observed in lymphomoids was consistent with patient’s response in the clinic. Lymphomoids are an innovative tool to assess treatment efficacy in clinically relevant contexts and could be used to uncover novel aspects of lymphoma biology.

## Introduction

Each tumor is characterized by a unique combination of molecular alterations and it establishes heterogeneous interactions with immune and stromal cell^1^. Tumor heterogeneity and plasticity hinder the selection of appropriate therapies for each cancer patient. To address this problem, experimental models based on patients’ biopsies have started to emerge and they are typically referred to as *patient avatars*^2^. These models are used to assess sensitivity to different therapies directly on tumor tissues derived from each patient to guide the selection of personalized treatments. In this context, patient-derived xenograft (PDX) models have been established in both solid tumors and hematological malignancies^3^. However, these models are time-consuming, lack the tumor microenvironment, and are more suitable to study aggressive tumors. As alternatives, the development of *ex vivo* models such as organoids and patient-derived tissue explants has allowed testing responses to therapy in some solid tumors ^4–11^. However, limited *ex vivo* models are available to anticipate therapeutic outcomes in B cell lymphomas. In particular, 3D models of B cell lymphomas are currently based on aggregates of available cell lines^12–15^, and spheroids^16–19^, which do not recapitulate the complex architecture of lymphoid tissue and are not representative of all lymphoma subtypes.

Non-Hodgkin lymphoma (NHL) is a diverse group of tumors originating from B cells at various stages of differentiation that accumulate heterogeneous genetic and epigenetic modifications^20–29^. NHL subtypes can be divided into indolent and aggressive tumors, where follicular lymphoma (FL) represents the most common subtype of indolent NHL, and diffuse large B cell lymphoma (DLBCL) the most common subtype of aggressive NHL^30–34^. Further subclassifications are based on the cell of origin^33^, molecular profile^35,36^, and their localization in distinct and specialized areas of the lymph nodes^37^. For example, FL, DLBCL, and Burkitt lymphoma (BL) originate from B cells in germinal centers, while mantle cell lymphoma (MCL) or marginal zone B cell lymphoma (MZL) arise from B cells in the mantle and marginal zones, respectively. In addition, the tumor architecture and the amount of interactions of lymphoma cells with the tumor microenvironment are variable among different tumor subtypes^38,39,40^. For instance, follicular lymphoma retains a follicular pattern, reminiscent of the lymph node’s architecture and cellularity, in contrast, DLBCL commonly exhibits architectural effacement^33^. Despite this heterogeneity, the first-line therapy is similar for most patients diagnosed with B cell lymphomas and it includes a combination of immunochemotherapy, where R-CHOP (Rituximab, cyclophosphamide, hydroxydaunorubicin, oncovin, prednisolone) is the most common regimen^41^. Between 60 and 70% of newly diagnosed B cell lymphomas are cured with R-CHOP treatment; however, patients with relapsed or refractory (R/R) disease have restricted therapeutic alternatives and their prognosis is limited, with low curative perspectives. As second-line treatments, T-cell-engaging therapies such as CAR-T cells and bispecific antibodies, and also small molecules blocking key oncogenic signaling pathways for the treatment of R/R aggressive lymphoma, FL, and MCL have been approved^42–45^. For instance, Bruton Tyrosine Kinase (BTK) inhibitors such as ibrutinib or acalabrutinib can be used for the treatment of SLL, MZL, and MCL, or lenalidomide has been approved for R/R FL, MZL, and MCL in combination with rituximab. However, the number of patients that will exhibit prolonged event-free survival remains low, possibly due to the remarkable heterogeneity of the disease. In this context, there is a need to develop reliable *ex vivo* models for lymphoma patients, where a large panel of treatments can be tested. To address this need, we first developed a strategy to culture *ex vivo* lymphoma tissue fragments retaining the architecture and cellular composition of the original tissue. We used this system using multiple human biopsies to test the effect of a panel of small molecule inhibitors. Selective sensitivity to the therapies was observed after a few days of treatment and for those cases where the *ex vivo* treatment matched the treatment of the patient in the clinic, it was predictive of the clinical outcome in 88.9% of the cases.

## Results

### Development of a system to maintain lymphoma explants *ex vivo*

B cell lymphoma arises in lymphoid organs which are composed of multiple immune cells and are organized in specialized areas. Thus, we wanted to develop a system that preserves the viability, proliferation, structure, and cellular composition of lymphoma tissues *ex vivo.* To this purpose, we adapted the air-liquid interface (ALI) method^46^ and we used this system to determine therapy efficacy in human lymphoma samples (**Figure 1A**). Initially, we optimized the conditions to maintain lymphoma explants in culture for several weeks by using secondary lymphoid organs isolated from vavP-Bcl2 transgenic mice. These animals develop tumors that recapitulate essential aspects of human FL^47^. We obtained tissue fragments between 0.75 to 1.5 mm^3^ from the spleens of aged vavP-Bcl2 mice that had developed FL. Tissue fragments were embedded in an RGD-hydrogel (arginyl-glycyl-aspartic acid hydrogel), which was diluted with different amounts of growth media to modulate the hydrogel stiffness (**Figure S1A)**. To sustain B cell viability, we added B cell activating factor (BAFF) (**Figure S1A)** and we tested it in combination with different cocktails of cytokines, chemokines, and small molecules. Flow cytometry analyses revealed that, except for BAFF, none of the additional molecules significantly improved viability, while IL2 significantly reduced the total number of B cell and germinal-center B cells after two weeks in culture (**Figure S1B-C, S2, and Table S1)**. Thus, we selected as optimal growth conditions intermediate hydrogel stiffness (1 part gel : 1 part growth medium) plus BAFF to guarantee good physical support to the tissue fragments while maintaining cell viability between 70-80% in the first week (**Figure S1A-C)**. Using these conditions, tumor fragments after one week largely retained the tissue architecture of the spleen with recognizable enlarged follicular structures (**Figure 1B**). Hence, we named these lymphoma explants *lymphomoids*. Next, we combined imaging and flow cytometry analyses to assess the tissue architecture and cellular organization of lymphomoids maintained in culture for four to five weeks (**Figure 1C-F**). By multicolor immunofluorescence staining, we were able to distinguish four main cell populations: B cells (B220), CD8+ T cells, CD4+ T cells, and macrophages (F4/80). After digitalization and quantification of the fluorescence signal intensity in multiple independent lymphomoids, we observed that the number of B cells was preserved *ex vivo*, although after three weeks CD4+ T cells tended to increase, while CD8+ T cells and macrophages gradually decreased (**Figure 1C-D**). Interestingly, the percentage of proliferating B cells was similar between lymphomoids and the original spleen from which the lymphomoids were generated, and it was maintained over time (**Figure 1E**). These results indicated that the cellular and structural composition of the original tissue was preserved for one to two weeks, and the proliferative capacity of the B cells for five weeks. Then, we used flow cytometry analyses to characterize B and T cell subpopulations. We quantified cell viability, the number of B cells (B220+), germinal center B cells (CD95+/GL7+), CD3+, CD8+, and CD4+ T cells, and CD4+ T-follicular helper cells (Tfh) for four weeks, and we used the original spleen from which the lymphomoids were generated as reference. First, we observed a reduction of 20-30% in overall cell viability in the first week in culture, which concerned primarily peripheral cells in the first 48h of the culture (**Figure 1F** and **Figure S1D-E)**. The number of B cells was maintained with a modest decrease after three weeks, similar to what was detected in the imaging analyses. This could be associated with a gradual loss of double-positive CD95+/GL7+ GC B cells (**Figure 1F**), although we noticed a tendency for B cells to lose CD95 expression, but not GL7 (**Figure S1F**). In addition, the percentage of CD3+, CD8+, and CD4+ T cells was constant, while CD4+ Tfh increased (**Figure 1F**). Hence, granular cytological analyses and imaging analyses were overall consistent. Lastly, to determine if the cells in the lymphomoids also retained the transcriptional profile of the cells in the original tissue, we analyzed six lymphomoids derived from the spleens of two animals, after one or two weeks in culture, and the original spleen tissues by single-cell RNA-sequencing. Cells clustered independently of the tissue of origin in two major groups: T cells identified by CD3 expression and B cells as CD79a positive (**Figure 1G-H**). Unbiased cluster analyses revealed the presence of two B cell subpopulations (cluster 1 and cluster 3), but not in T cells (cluster 2) (**Figure 1I**). Cluster 1 grouped B cells from the two original spleens and from lymphomoids maintained in culture for one week, while cluster 3 grouped B cells from lymphomoids in culture for two weeks (**Figure 1I and Figure S3A)**, suggesting that the expression profile of B cells in lymphomoids started to change after one week in culture. To better define different B and T cell subtypes in the original spleens and lymphomoids, we used a curated list of genes as markers to distinguish specific cell types (**Table S2**)^48^. Interestingly, the distribution of different cell populations was similar between the original tissues and lymphomoids maintained in culture for one week (**Figure S3A)**, indicating that they retained similar expression patterns and cellular composition. Conversely, after two weeks in culture, lymphomoids were enriched in B cells expressing plasma cell markers (**Figure S3B**). Indeed, differential expression analyses between B cells from lymphomoids in culture for one or two weeks showed upregulation of late B cell differentiation markers such as *Xbp1* and *Zbp1*^49,50^, and downregulation of early B cells differentiation markers such as *Ebf1*, *S1pr1*, and *Pou2f2*^51–53^ (**Figure S3C**). However, despite this initial transcriptional shift, we did not detect major changes in the expression of differentiation markers on the B cell surface (e.g., CD138+ cells) (**Figure S3D**). Overall, morphological, cytological, and molecular characterizations indicate that lymphomoids retain the features of the original tissue for up to one week in culture, defining this as the ideal time frame to test sensitivity to therapies in this model.

**Figure 1:**
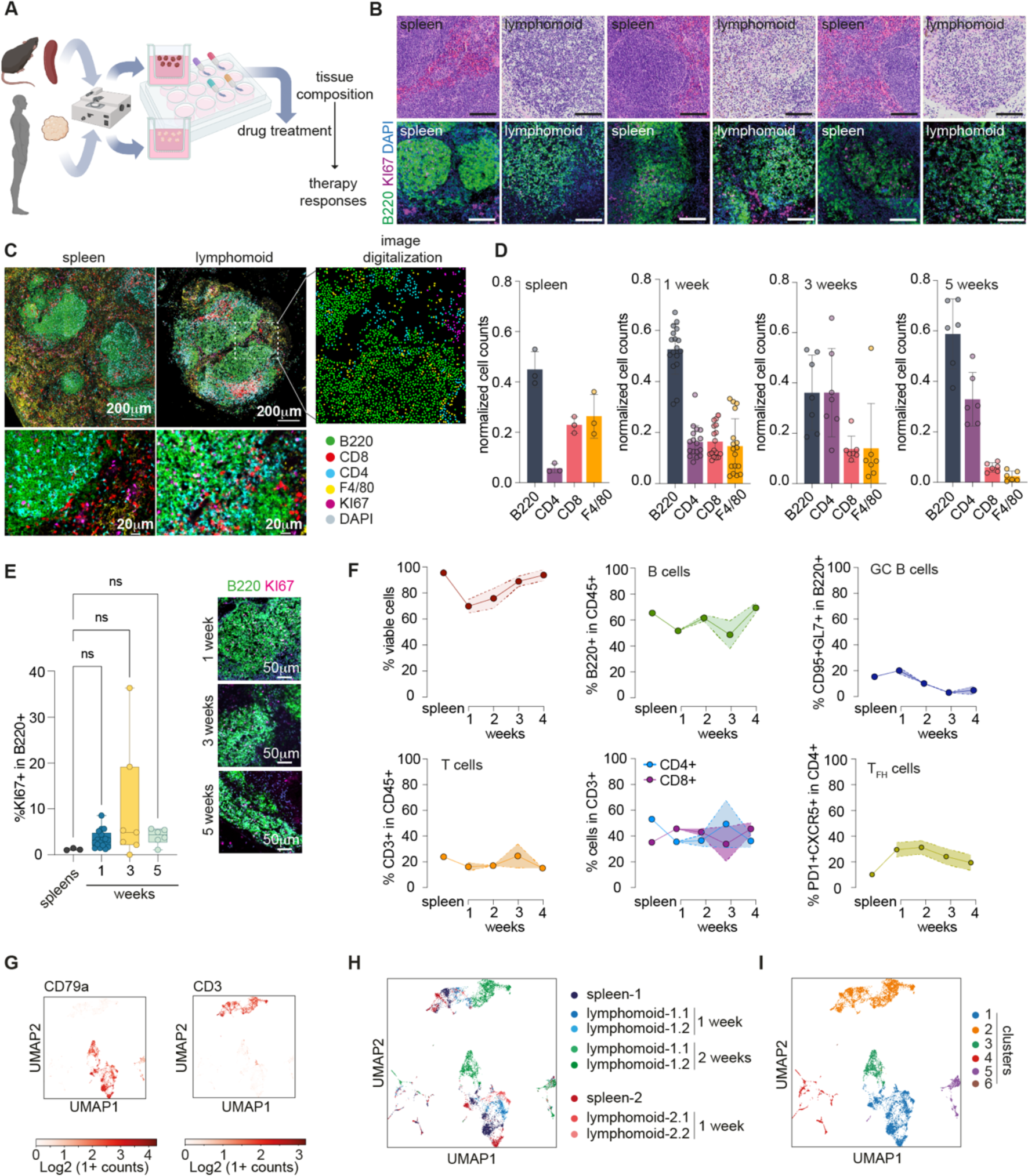
A system to maintain lymphoma biopsy *ex vivo*. (A) Schematic of the procedure to process and test sensitivity to therapies for murine and human samples (created with BioRender.com). (B) Hematoxylin and eosin staining of a vavP-Bcl2 mouse spleen and a representative lymphomoid derived from the same spleen (upper images), and B220+KI67 immunofluorescence of vavP-Bcl2 spleens and lymphomoids. scale bar 100 μm. (C) Representative images of 5 color immunofluorescence staining for the indicated markers of a vavP-Bcl2 mouse spleen and a lymphomoid derived from the same spleen at two different resolution and a digitalized image for the lymphomoid as a magnified cropped detail. (D) Quantification of different cell types based on fluorescence signal at different time points (the data points represent different lymphomoids and sections from different vavP-Bcl2 mice; spleen n=3, 1 week n=17, 3 weeks n=7, 5 weeks n=6). (E) Quantification of fluorescence signal to detect proliferating B cells at different time points (spleen n=3, 1 week n=17, 3 weeks n=7, 5 weeks n=6). Data is shown in box and whiskers (min to max) with all data points. (F) Quantification of different cell types by flow cytometry of vavP-Bcl2 mouse lymphomoids at different time points (n=4 lymphomoids for each time point). (G-I) UMAP projection of single cells RNA-sequencing cells color coded for cells expressing CD79a or CD3 genes (G), color coded by the origin of the samples (vavP-Bcl2 spleen n=2, lymphomoids 1 week (n=4), and lymphomoids 2 weeks (n=2) (H), color coded based on the different cell clusters found in the analysis.

### Human lymphomoids retain the cell heterogeneity of lymphoma tissues

To assess if lymphomoids could be used to test sensitivity to therapies, we generated and characterized lymphomoids from fresh human biopsies obtained from 25 patients with suspected lymphoma. Part of the tissues was rapidly processed to create lymphomoids and the remaining tissue was fixed and used for comparative analyses. We retrospectively retrieved available clinical information for these patients confirming lymphoma diagnosis, subtype classification, and genomic alterations (**Table S3**). This cohort included nine diffuse large B cell lymphoma, five primary mediastinal B cell lymphoma, two high-grade B cell lymphoma, two small lymphocytic lymphoma, one mantle cell lymphoma, one marginal zone lymphoma, two follicular lymphoma, one transformed follicular lymphoma, one Burkitt lymphoma, and one T cell lymphoma (**Figure 2A**). The lymphomoids obtained from these biopsies were maintained in culture for three to seven days, and for each patient they were treated with different small molecule inhibitors (**Figure 2A**). First, we assessed the morphological quality of lymphomoids by histopathology analyses (hematoxylin and eosin, H&E staining) compared to the original tissue. Initial analyses revealed that nineteen biopsies were of high quality, three samples suffered from mechanical trauma during the surgical procedure or were poorly preserved, two samples were mainly fibrotic (patient 4 and 7) and the lymphomoids obtained from one sample were rapidly contaminated with fungi (patient 5), probably because the biopsy was obtained through the oral cavity (**Figure S4A)**. Among the nineteen high-quality tissues, lymphoma cells could be detected in seventeen samples, while two samples contained mostly reactive lymph node tissue (**Figure S4B**). We analyzed the cell composition of lymphomoids originating from these tissues over multiple sections by multicolor fluorescent imaging using CD20 for B cells, CD8 and CD4 to distinguish T cells, CD68 for monocytes, and Ki67 for proliferation (**Figure 2A-B** and **Figure S4C**). We compared the proportions of cells in the different subpopulations in multiple untreated lymphomoids and the original tissue and we observed a significant correlation for each cell type (**Figure 2c and Figure S4D**). In addition, we computed a similarity index considering all cell types together (non-proliferating CD20+ B cells, proliferating CD20+ B cells, CD4+ T cells, CD8+ T cells, CD68+ macrophages, other cells) for each tumor sample and lymphomoid. In general, lymphomoids are significantly more similar to their original tumor than to other tumors, and lymphomoids from the same tumor are more similar to each other than to lymphomoids originated from another tumor (**Figure 2D**). Importantly, lymphomoids from the same tumor are as similar among each other as they are with the tumor from which they were derived. To further characterize the spatial and molecular structure of human lymphomoids and their tissue of origin, we performed spatial transcriptomics analyses on original tissues and lymphomoids obtained from patient 8 and patient 3 using the 10x Genomics Visium technology. The original tissue obtained from patient 8 showed that lymphoma cells were ubiquitously spread in the tissue and mixed with other cell types (**Figure 2E**). In addition, we distinguished areas enriched with densely clustered lymphoma cells (red line), stroma (green line), and blood vessels (blue line) (**Figure 2E**). The tissue obtained from patient 3 contained reactive follicles surrounded by areas with sinus histiocytosis and blood vessels, but not lymphoma (**Figure S5A)**. Spatial transcriptomics analyses allowed to assay the full transcriptome in multiple ‘spots’ regularly interspersed across the tissue, each containing 20-25 cells (**Figure S5bB-C**).

**Figure 2:**
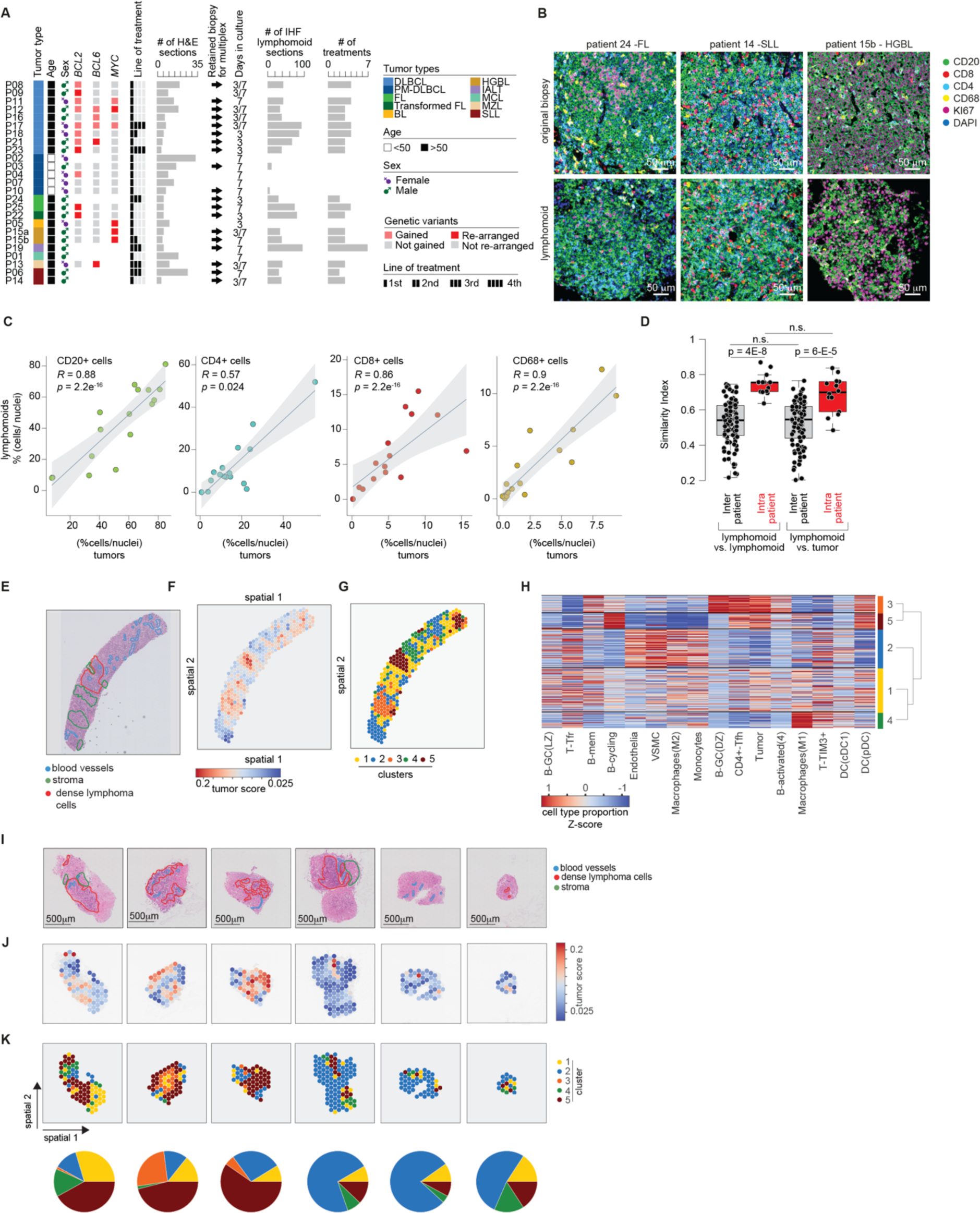
Characterization of human lymphomoids obtained from fresh tumor biopsies. (A) Graphical summary of all the cases collected in this study. (B) Representative images of multicolor immunofluorescence staining of indicated markers of three human lymphoma biopsies (upper panel) and their respective lymphomoids (lower panel). Scale bar 50 μm. (C) Spearman’s correlation coefficient analyses of untreated lymphomoids vs original tumor biopsies on the different markers analysed by multiplex IHF (n=15 tumors; one tumor was excluded from this analysis since it was CD20 negative). The data is shown as a percentage and was normalized using weighted ratios to the total number of nuclei detected in each section. (D) Similarity index comparisons in the indicated conditions. (E) H&E staining of patient 8 DLBCL biopsy. Tissue features are highlighted with lines of different colors. (F-G) Representation of 10X Genomics Visium spots color-coded by the tumor score (F) and based on unbiased clustering (g). (H) Heatmap of cell type enrichment in each spot of the biopsy determined using Bayesprism deconvolution. Only top 4 differentially enriched cell types in each cluster are shown. The cell state proportions are z-scale normalized for each cell type. The dendrogram shows the distances between the different clusters. (I) H&E staining of six lymphomoids maintained in culture for 3 days. Tissue features are highlighted with lines of different colors. Scale bar 500 μm. (J-K) Representation of 10X Genomics Visium spots color-coded by the tumor score (J), based on cluster scores obtained in the original tissue (K-top), and pie-charts summarizing the cluster composition of each lymphomoid (K-bottom).

To overcome the lack of single-cell resolution, we used the BayesPrism method to predict the cell type composition of each spot and deconvolve spatial transcriptomic data into cell type-specific mRNA expression matrices^54^. To this purpose, we generated a reference dataset using single-cell RNA-seq data from 37 normal cell types and subtypes obtained from secondary lymphoid organs^55^ and one DLBCL patient^56^. Then, we applied BayesPrism to predict the proportions of these 38 cell types in each spot (**Table S4**). Importantly, with this approach, we could identify multiple spots with a high proportion of tumor cells in patient 8, but not in patient 3, in agreement with the histological classification (**Figure 2F** and **Figure S5D**). Next, we unbiasedly clustered the spots based on their cell type proportions, independently on patient 8 and patient 3, and identified 5 clusters in patient 8 and 6 clusters in patient 3 (**Figure 2G** and **Figure S5E**). In patient 8, the spots in clusters 3 and 5 were enriched in characteristic lymphoma cell types: proliferating B cells, CD4+ Tfh cells, and GC B cells. Notably, cluster 5 overlapped with the area enriched for lymphoma cells identified by histological analyses (**Figure 2E, G**). Spots in clusters 2 and 4 were predominantly composed of monocytes, macrophages, and endothelial cells, while cluster 1 comprised spots with mixed cell type composition (**Figure 2H**). In patient 3, areas annotated with reactive lymph nodes by histology were grouped in one cluster (cluster 4), which included GC B cells, CD4 Tfh cells, and follicular dendritic cells; spots in cluster 2 were mainly composed of macrophages, endothelial cells, and T follicular reticular cells; cluster 0 included different groups of activated B cells and B cells with high expression of interferon-induced genes (B-IFN); and spots in cluster 1 were enriched in proliferating B cells, plasma cells, and tumor cells, although the tumor score in this sample was low (**Extended Data Figure 5F**). Thus, this spatial transcriptomics analysis led to the identification of spatial cell niches, which matched with the histological assessment of the tissue. Next, to determine if the same niches were present in six different lymphomoids derived from patient 8 maintained in culture for three days, we deconvolved the expression profile of each spot using BayesPrism, and based on their cell type proportion they were assigned to one of the 5 clusters determined in the original tissue. Interestingly, each lymphomoid preserved the heterogeneity of the original tissue with spots enriched for tumor cells being observed in all lymphomoids (**Figure 2J**) and all 5 clusters being recapitulated in different proportions (**Figure 2K**). Overall, both multicolor imaging and spatial transcriptomics analyses confirmed that it is possible to maintain human lymphomoids in culture for a few days, while preserving the heterogeneity of the original tissue composition and the proliferation of tumor cells.

### Human lymphomoids can be used to test drug sensitivity in lymphoma patients

Different therapies have been approved as second-line treatments for lymphoma patients. However, there are no knowledge-based methods to select the best treatment for each patient. Hence, we assessed the possibility of using lymphomoids to screen in parallel multiple small molecule inhibitors and evaluate patient-specific tumor sensitivities. Multiple lymphomoids obtained from 14 different patients (6, 8, 11-14, 16-18, 21-25) were treated with ibrutinib (BTK inhibitor), idelalisib (PI3Kδ inhibitor), lenalidomide (E3 ubiquitin ligase modulator), venetoclax (BCL2 inhibitor), or tazemetostat (EZH2 inhibitor). Lymphomoids from two independent and sequential biopsies (LP15A and LP15B) of an early relapsed case of high-grade B cell lymphoma were treated with alisertib (Aurora kinase A inhibitor), everolimus (mTOR inhibitor), and lenalidomide. Lastly, lymphomoids from a T cell lymphoma were treated with panobinostat (HDAC inhibitor), 5-azacytidine (methyltransferase inhibitor), tazemetostat, idelalisib, alisertib, lenalidomide, and venetoclax. The efficacy of these treatments was determined based on the difference of proliferating B cells (fold-change) that was detected in treated vs. untreated samples (**Figure 3A-B**). This analysis revealed heterogeneous responses to the same or different compounds among patients (**Figure 3B**). For instance, lymphomoids derived from patient 8 were exclusively sensitive to ibrutinib, while lymphomoids derived from patient 21 were only sensitive to idelalisib, and those from patients 12, 14, 17, 18 showed similar sensitivity to both ibrutinib and idelalisib (**Figure 3B**). Additionally, to distinguish between cytotoxic and cytostatic effect, we computed the fold-change of total B cells between treated and untreated conditions (**Figure 3C**). In general, we observed lower fold-changes for proliferating B cells compared to the total number of B cells, suggesting that mainly B cell proliferation was affected by these treatments (**Figure 3B-C**). Nonetheless, we also observed a concomitant reduction of total and proliferating B cells upon treatment with ibrutinib, idelasilib, and venetoclax in multiple patients, suggesting that these drugs may also exert cytotoxic effects. For patient 15, we obtained two sequential biopsies taken 7 months apart, before and after tumor relapse (LP15A and LP15B, respectively). Lymphomoids derived from this patient were treated with the same compounds and showed similar sensitivity to everolimus and alisertib, but not to lenalidomide. Interestingly, whereas in the first biopsy (L15A) the effect was largely cytostatic, in the second biopsy (L15B) we observed cytotoxic effects (**Figure 3B-C** and **Figure S6A-B**). We also compared the effect of three or seven days of treatment on lymphomoids derived from 5 different patients, but we did not observe major differences (**Figure S7**). Next, we used spatial transcriptomics analyses to assess sensitivity or resistance to small molecule inhibitors based on the estimated proportion of tumor cells, and to understand the effect of these therapies not only on tumor cells but also on the tumor microenvironment composition. The proportion of tumor cells in lymphomoids obtained from patient 8 treated with ibrutinib and idelalisib confirmed sensitivity to ibrutinib but not idelalisib (**Figure 3D**). Indeed, clustering of spots based on gene expression in untreated and treated lymphomoids showed that ibrutinib-treated lymphomoids formed a distinct cluster (cluster 4) while clusters 1 and 2 were composed of both untreated and idelalisib-treated samples (**Figure 3E**). Importantly, in ibrutinib-treated lymphomoids, we noticed upregulation of several inflammatory genes (e.g., *IFI27*, *IFI6*, *IFIT3*) and loss of expression of B cell markers (**Figure 3F**), indicating that ibrutinib affected viability and proliferation of B cells and triggered the activation of inflammatory anti-tumoral responses. Next, to decipher if these compounds affected different subpopulations of the tumor microenvironment, we compared the proportion of different cell populations in untreated and treated lymphomoids. In lymphomoids treated with ibrutinib, we detected a strong reduction of tumor cells, GC B cells, and proliferating B cells, and an increase of CD8+ cytotoxic T cells and macrophages (**Figure 3G** and **Table S5**). Conversely, in the idelalisib-treated lymphomoids the proportion of tumor cells was largely unchanged, but GC-B cells, CD4+ Tfh cells, and FDC were enriched, and proliferating and activated B cells were depleted (**Figure 3G**), indicating that treatment with idelalisib, although not effective on tumor cells, was influencing normal B cells. Hence, lymphomoids represent a valid method to test in parallel the activity of different therapies and, in combination with spatial analyses of the tissue, they could reveal how the tumor microenvironment reacts to drug treatment.

**Figure 3:**
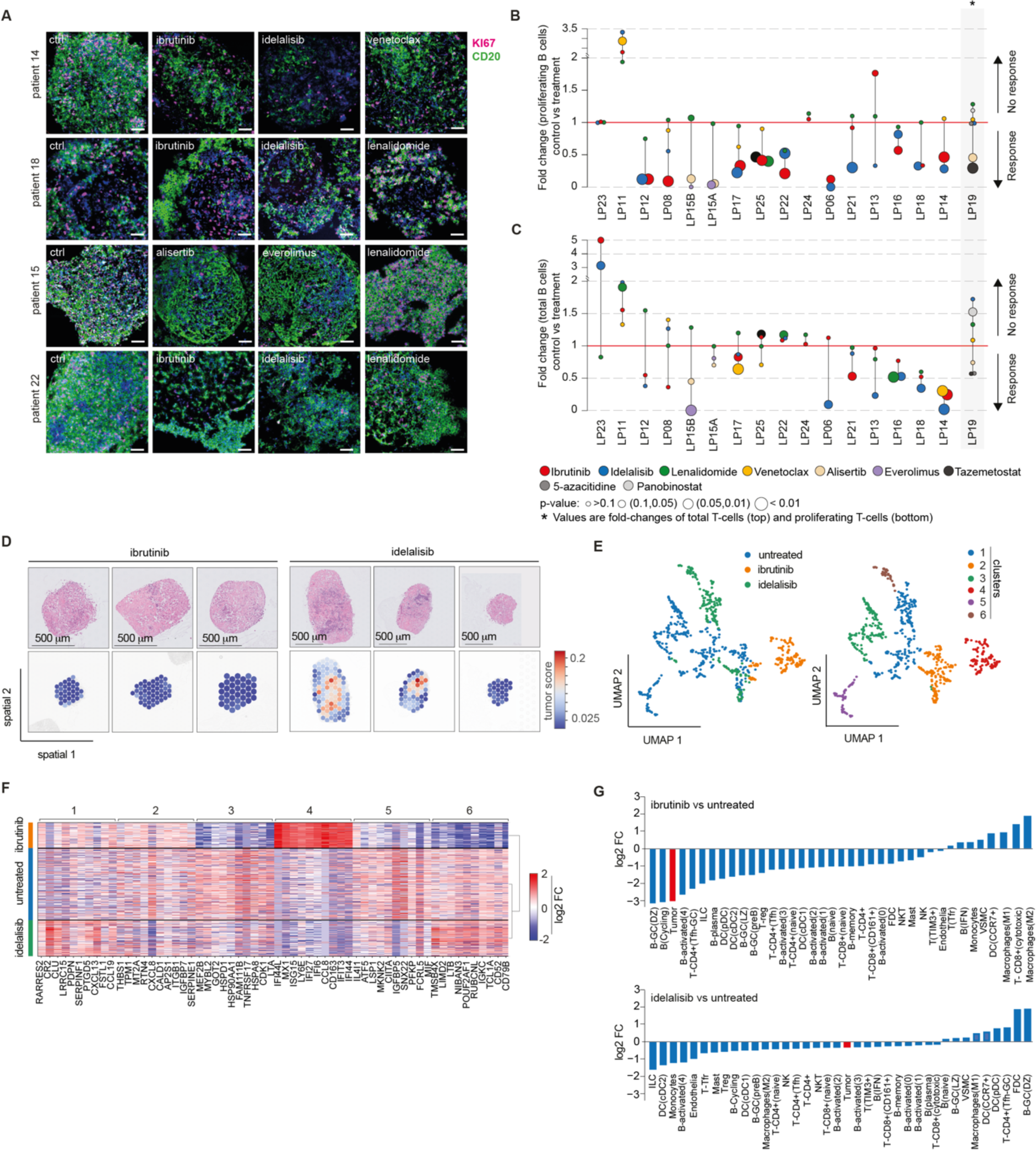
Sensitivity to targeted therapies using patient-derived lymphomoids. (**A)** Representative images of immunofluorescence staining for KI67 and CD20 in lymphomoids derived from 4 different biopsies. Scale bar 50 μm. **b-c**) Proportion of proliferating (B) and total number (C) of cells compared to controls. Each treatment is color-coded and the p-values were calculated using non-parametric ANOVA followed by Dunn’s test. (D) H&E and tumor score of ibrutinib and idelalisib treated lymphomoids. (E) UMAP projection of 10X Genomics Visium spots colored by sample (left) and by cluster (right). (F) Heatmap of differentially expressed genes in each cluster and sample obtained in the 10X Genomics Visium experiment. (G) Comparison of cell type enrichment in ibrutinib (top) and idelalisib (bottom) treated lymphomoids versus control.

### Lymphomoids to anticipate clinical responses

To assess the predictive potential of our lymphomoids platform, we retrieved treatment response data for 6 patients, among our donors, which were treated with the same compounds that we tested in lymphomoids derived from their original tumor. Two patients diagnosed with SLL (patient 6 and patient 14) were treated with either a combination of obinutuzumab (anti-CD20 antibody) and acalabrutinib (BTK inhibitor), or acalabrutinib alone (**Figure 4A-B**). In patient 6- and patient 14-derived lymphomoids (LP06 and LP14, respectively), we observed a significant reduction of proliferating B cells when treated with ibrutinib, with complete depletion of proliferating B cells in LP06 (**Figure 4A**) and a residual 10% of Ki67+ B cells in LP14 (**Figure 4B**). Consistently, in the clinic these patients showed either complete (patient 6) or partial (patient 14) response (**Figure 4a-b**). Patient 11 was diagnosed with DLBCL and treated with rituximab (anti-CD20 antibody) and revlimid (lenalidomide) as an alternative to chemotherapy. Lymphomoids obtained from the tumor biopsy showed that the tumor was refractory to lenalidomide (**Figure 4C**). Consistently, after 2 cycles of treatment, radiological assessment by PET-CT showed no metabolic response (NMR) with an overall disease progression (Deauville score 5) for patient 11 (**Figure 4C**). A second patient diagnosed with R/R DLBCL (patient 17) was initially treated with ibrutinib and subsequently with tafasitamab (anti-CD19) and lenalidomide. Both treatments resulted in NMR with a Deauville score of 5 (**Figure 4D**). Lymphomoids derived from this patient (LP17) were treated with lenalidomide and ibrutinib. Although they exhibited sensitivity to ibrutinib, they confirmed the resistance to lenalidomide (**Figure 4D**). Patient 18, another patient with R/R DLBCL was treated with ibrutinib and at the baseline PET-CT showed a complete metabolic response (CMR) following three weeks of ibrutinib monotherapy. Similarly, ibrutinib led to a significant reduction of proliferating B cells, confirming sensitivity to this molecule (**Figure 4E**). Patient 13 was diagnosed with relapsed MZL and was treated with ibrutinib, but the treatment was interrupted after three weeks due to severe bleeding, and the PET-CT showed tumor progression. Thus, the treatment was switched to a combination of tafasitamab and lenalidomide, but the tumor did not respond (**Figure 4F**). Because ibrutinib had only been given for a short period and had demonstrated high efficacy in clinical trials with MZL patients^57^, patient 11 was treated again with ibrutinib, but the tumor remained refractory to this treatment as confirmed by PET-CT with NMR and a Deauville score of 5 (**Figure 4F**). Lymphomoids obtained from the relapsed tumor (LP13) were treated with ibrutinib and lenalidomide, but neither compound was effective, consistent with the observed clinical response. Finally, patient 24 was diagnosed with relapsed follicular lymphoma and treated (3rd line) with tafasitamab and lenalidomide, but after two cycles the PET-CT revealed that the lesions did not respond to the treatment (**Figure 4G**). Likewise, lymphomoids from the same patients did not exhibit sensitivity to lenalidomide. Overall, sensitivity and resistance to the treatments measured in lymphomoids largely matched the patient’s response (8/9 tested conditions, 88.9%), indicating that lymphomoids could provide valuable information to guide therapeutic choices in the clinic.

**Figure 4:**
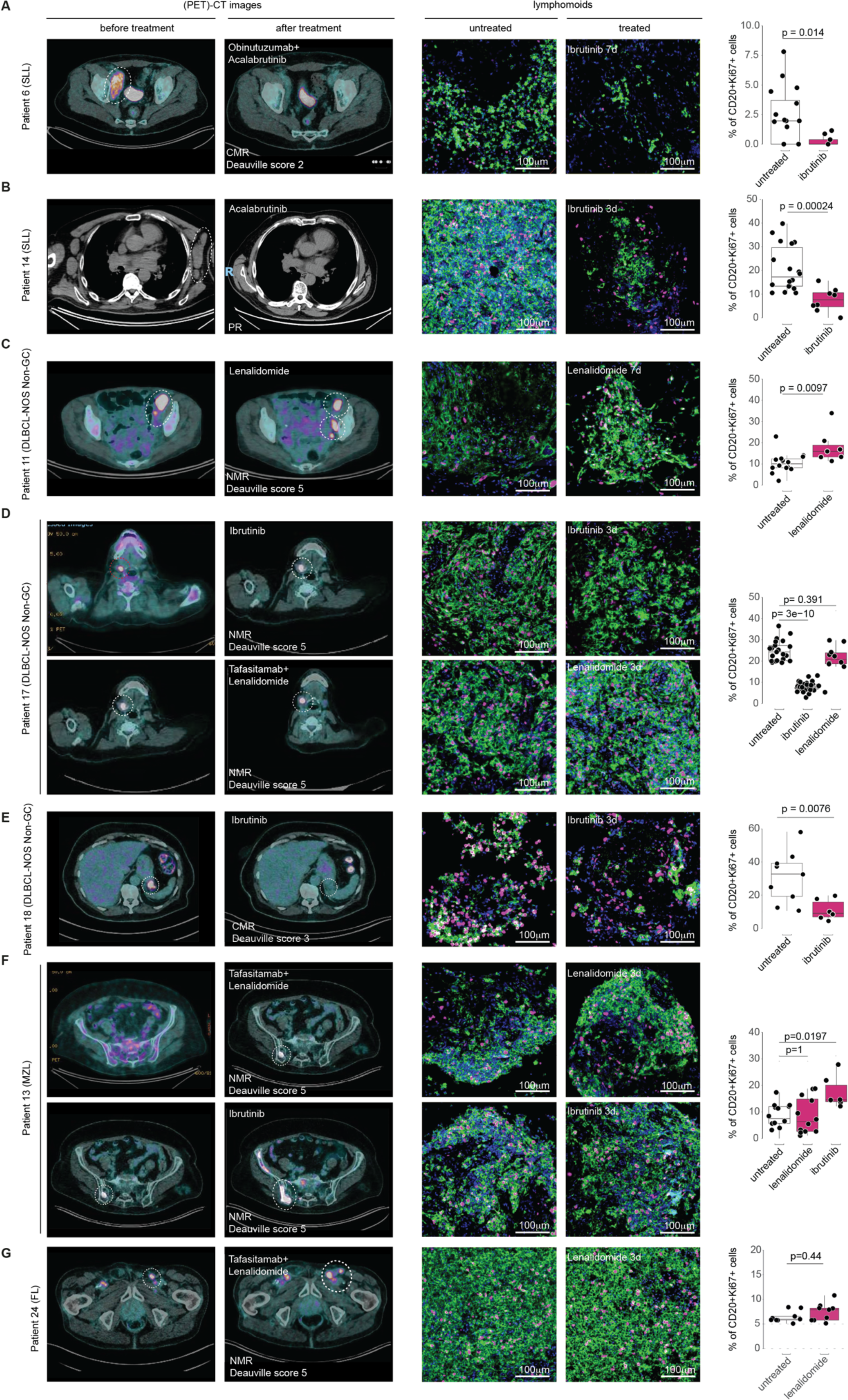
Matched results obtained in the clinic and on lymphomoids in seven patients. (A-F) PET-CT before and after treatment, representative image and quantification (of the CD20 and KI67 immunofluorescence signal of the lymphomoids derived from patient 6 (A), patient 14 (B), patient 11 (C), patient 17 (D), patient 18 (E), patient 13 (F), and patient 24 (G). Data was analyzed by either Wilcoxon rank-sum test or Kruskal-Wallis followed by Dunn’s posthoc test. CMR: complete metabolic response; NMR: no metabolic response; PR: partial response.

## Discussion

The ultimate goal of personalized medicine approaches is to tailor medical treatment to individual patients. However, to fulfill this promise, predictive models of each tumor are required to assess treatment response. Ideally, to be routinely used in the clinic, these models should be suitable to test multiple conditions and have simple but reliable markers to rapidly determine sensitivity or resistance to therapies. The culture of *ex vivo* tissue fragments obtained from patient biopsies represents an ideal system to test different therapies in a clinically reasonable time frame. For example, tissue biopsies from a number of solid tumors have been used to measure sensitivity to immunotherapies in 48h^5^. In lymphoma, human cell lines and patient-derived animal models have been extensively used for preclinical studies^56,58–60^. However, these models are not suitable to study treatment sensitivity in indolent disease and they do not preserve the tissue composition and its spatial organization. Lymphomoids generated from intact tissue fragments overcome these limitations and could represent a reliable alternative to test sensitivity to different therapies in a clinically actionable time. In this context, measuring proliferating B cells could be sufficient to select the most effective therapy for each patient. Interestingly, the analysis of lymphomoids from two independent biopsies taken 7 months apart on a high-grade B cell lymphoma patient that relapsed after being treated with chemotherapy revealed that they were still sensitive to the same compounds. These results suggest that in early relapsed cases, the lymphomoid technology could also be used to anticipate response to therapies of the relapse tumor, in agreement with a recent study showing that refractory or early (<12 months) relapsed disease share similar mutation profiles and treatment outcomes as the first lesion^61^. However, to fully determine the clinical impact of our approach, prospective trials with matching therapies need to be designed. Ideally, lymphomoids should be used in clinical trials where patients will be enrolled based on specific inclusion criteria, and the clinical endpoint is predefined. Matching clinical responses and lymphomoid results would consolidate the validity of this method in the clinic. In addition, the automatization of tissue handling, slide preparation, and imaging data analyses will be critical to use this system for routine diagnostics. Nevertheless, the concordant results that we obtained in seven matching cases suggest that this approach could be clinically relevant and could be used to uncover features in the tumor tissue architecture that dictate therapy sensitivity or resistance.

In parallel to their clinical and preclinical application, the lymphomoids could serve as a system to further understand tissue remodeling after treatment using spatially resolved molecular assays. Indeed, the initial analyses that we performed revealed rearrangements in the tissue organization and immune cell subpopulations upon treatment. Interestingly, we noticed that treatment with ibrutinib induced an inflammatory response and enrichment of activated CD8+ T cells, macrophages, and dendritic cells (CCR7+). This opens the possibility of directly studying immune cell activation and their functional interdependency using lymphomoids. In addition, it is possible to envision that lymphomoids could be generated from normal lymph nodes and they could serve as an alternative method to the self-organizing lymphoid tissue system that has been recently proposed^62^. Overall, lymphomoids represent both a new experimental system to study biological processes in B cell malignancies and a new technology to model individual patient tumors and inform treatment decisions in the clinic.

## Methods

### KEY RESOURCES TABLE

### RESOURCE AVAILABILITY

#### Lead contacts

Further information and requests for resources and reagents should be directed to and will be fulfilled by the lead contacts, Elisa Oricchio and Albert Santamaria-Martínez.

#### Materials availability

The mouse line used in this study (vavP-Bcl2) was obtained from Prof. Suzanne Cory.

### EXPERIMENTAL MODEL AND STUDY PARTICIPANT DETAILS

#### Human sample collection

Patients with a clinical suspicion of abdominal or axillary lymphoma and scheduled for a surgical or needle lymph node biopsy were informed about the project and offered to participate in the study. Upon informed consent, fresh biopsies from lymphoid tissue were collected in 50 ml Falcon tubes containing 30mL of cold PBS + 1x normocin. The samples were then immediately transported on ice inside a polystyrene box to the lab, where they were processed as described below. All procedures were carried in accordance with the Ordinance on human research with the exception of Clinical trials (HRO) and were previously approved by the *Commission cantonale d’éthique de la recherche sur l’être humain* (CER-VD, project-ID 2020-02532).

#### Animal models

vavP-Bcl2 (C57BL/6) mice were bred and housed in ventilated cages in the conventional mouse husbandry of EPFL. All the experiments were carried out in accordance with the Swiss Animal Welfare Regulations and were previously approved by the Cantonal Veterinary Service of the Canton de Vaud (license VD3631).

### METHOD DETAILS

#### Tissue culture

Tumor tissues were placed in PBS and rapidly processed with a McIlwain Tissue Chopper (Ted Pella Inc.) producing fragments of 0.75-1.5mm^3^. Four to ten explants were then placed in inserts with permeable, membranous bottom (Millicell-CM, Millipore) containing a 1:1 solution of Vitrogel RGD High Concentration (TheWell Bioscience) and RPMI1640 supplemented with 5% FBS, 1x normocin, 1x insulin/selenium/transferrin, and 1x sodium pyruvate. The insert was placed in a 12 well plate and left in a 5% CO_2_ incubator at 37°C for 20 minutes for the matrix to gelify. Finally, RMPI1640 complete + 0.2μg/ml hBAFF (Biolegend) was added to each well. Media was replaced every 3-4 days.

#### Immunostaining

Formalin-fixed paraffin-embedded (FFPE) tissues were sectioned at 5 mm in a Hyrax M25 microtome and placed in Superfrost^TM^ slides. 5-plex immunohistofluorescent staining was performed on the fully automated Ventana Discovery Ultra (Roche Diagnostics) using the manufacturer’s solutions and antibodies. Briefly, deparaffinized and rehydrated FFPE sections were pretreated with heat using standard conditions (40 minutes) in CC1 solution. Primary antibodies (human): CD4 (SP35), CD8 (SP57), CD20 (L26), CD68 (KP-1), KI-67 (30-9), all of them ready-to-use. Primary antibodies (mouse): B220, CD4, CD8, F4/80. The images were acquired in an Olympus VS120 whole slide scanner and analyzed using QuPath.

#### Image digitalization and analysis

The image analysis pipeline is based on QuPath^63^, DeepCell ^64^, MCMICRO ^65^, and custom scripts implemented in Groovy, Python and RSource code. Briefly, .vsi images acquired on the Slide Scanner (Olympus), were opened as a project on QuPath. First, lymphoid boundaries were drawn and exported using a custom script. Then, the fluorescence thresholds for each channel were calibrated in each image and exported using a second script. Next, the images were converted to .ome.tiff and DeepCell was run to detect the nuclei and the cytoplasmic areas, which were used for marker quantification of the nuclear and cytoplasmic markers, respectively. Marker quantification was performed using MCMICRO by taking, for each cell, the mean intensity of each marker within the respective segmentation mask. Finally, a cell was classified as “positive” for a given marker if the difference between the cytoplasmic intensity of that marker and its calibrated threshold was the highest across markers. Cells with intensities below all marker thresholds are deemed as coming from an unknown cell type and called “otherCell”. Additionally, cells assigned to markers are further classified as proliferating or not according to whether their nuclear Ki67 intensity is above or below its calibrated threshold. Lymphomoid-level cell composition is obtained by intersecting the spatial coordinates of classified cells with the lymphomoid boundaries saved at the first step.

#### Flow cytometry

Spleens, lymph nodes, and lymphomoids were mechanically disaggregated, filtered through a 0.40mM nylon mesh strainer and resuspended in cold FACS buffer (2%FBS, 1mM EDTA in PBS). Erythrocytes were lysed using a Red Blood Cell Lysis buffer (Biolegend) for 2 minutes at room temperature and then cells were washed in PBS and incubated for 15 minutes at room temperature with a PBS solution containing the fixable viability dye Zombie Violet^TM^ (Biolegend, 1:100) and CD16/CD32 Fc Block (BD Biosciences, 1:100). Next, cells were washed and stained with the following fluorescent-labeled antibodies for 30 minutes at 4°C: Biolegend: CD3e-FITC (145-2-C11, 1:300), CD4-AF700 (RM4-5, 1:300), CD8-BV570 (53-6.7, 1:200), CXCR5-PerCP-Cy5 (L138D7, 1:300), PD1-PECy7 (29F.1A12, 1:300), CD95-PE (SA367H8, 1:200), CD138-PE-Cy7 (281-1, 1:300); BD Pharmingen: CD45-APCCy7 (30-F11, 1:500), T- and B-cell activation antigen-AF647 (GL7, 1:200); BD Horizon: B220-PECF594 (RA3-6B2, 1:500). Finally, cells were washed, fixed in BD Cytofix/Cytoperm Solution (BD Biosciences), washed again and analyzed in a Gallios machine (Beckman Coulter). Data was analyzed using FlowJo 10.8 (Becton Dickinson).

#### Single-cell RNA sequencing and data processing

vavP-Bcl2 spleens and lymphomoids were mechanically disaggregated, filtered through a 0.40mM nylon mesh strainer and resuspended in PBS. Erythrocytes were lysed using a Red Blood Cell Lysis buffer (Biolegend) for 2 minutes at room temperature and then cells were washed in PBS, resuspended in medium and processed for sequencing using the SingleCell 3’ Reagent Kit v3.1 and a 10x Chromium single cell controller. The sequencing depth was between 183 to 225 million reads/sample. Raw sequencing reads were aligned to mouse reference genome GRCm38 using 10x Cell Ranger pipeline v6.0.2. Count matrices were further processed in Python v3.9.1 using Scanpy v1.9.1 toolkit. Cells with less than 200 genes expressed, and genes detected in less than 3 cells were removed. For the first murine tumor and lymphomoids, cells with total counts between 2000 and 75000 expressed genes between 800 and 5000, and counts from mitochondrial genes under 10% were retained. For the second tumor, cells with total counts between 2000 and 100000, counts from mitochondrial genes under 15%, and between 1000 and 8000 total genes expressed were shortlisted. The two parental tumor samples and corresponding lymphomoids counts matrices were concatenated. Merged counts were normalized for total 100,000 counts per cell, and log transformed. Top 2000 highly variable genes were used to construct a UMAP projection of the cells with top 50 principal components, 15 neighbors and 0.2 as the effective minimum distance (min_dist parameter). Finally, Leiden clustering algorithm was used with 0.1 resolution to obtain 6 clusters.

#### Spatial transcriptomics

##### Tissue and slide preparation

Formalin-fixed paraffin-embedded (FFPE) tissues were sectioned at 5 mm in a Hyrax M25 microtome and placed in a Visium slide. The slides were then deparaffinized, stained with H&E, and images were acquired using an Olympus VS120 whole slide scanner. Next, sections were decrosslinked, and were subsequently treated as per manufacturer’s instructions to hybridize the probes, ligate them, and finally generate FFPE libraries. Samples were sequenced using Illumina’s NextSeq.

#### Spatial Transcriptomics data preprocessing

For each capture area in the 10x Visium slide, FASTQ files and H&E images were processed using Space Ranger v1.3.0. Staining images were aligned and spots covered by tissue were selected using Manual Alignment for Space Ranger wizard found in the Loupe Browser software v5.1.0. The aligned and selected spots were passed to Space Ranger call using the loupe-alignment argument. The raw reads were aligned to hg38 human reference genome (refdata-gex-GRCh38-2020-a). The aligned reads were analyzed using Python v3.9.1 using Scanpy v1.9.1 toolkit. A median of 26062 counts was obtained, with 7080 median genes per spot.

#### Cell detection in Visium slides H&E images

For each capture area of the 10x Visium slide, the corresponding H&E images were processed using the inbuilt cell detection option in QuPath v0.3.2 with default parameters. The detected cell measurements (centroid, area etc.) were exported to a tab-separated file, and an in-house R script was used to compute the number of cells identified within each visium spot.

#### 10x Visium data deconvolution and processing

Count values for each spot of the 10x Visium slide was deconvolved using BayesPrism v2.0, using the single-cell reference dataset of the 38 cell types. The “final” theta values from the BayesPrism fitted object were for downstream analysis. Parental Visium spots’ deconvolved signals were used to cluster the spots using Leiden clustering algorithm at 0.2 resolution. Marker cell types and marker genes for the clusters were calculated using Scanpy’s rank_gene_groups function, with Wilcoxon test method.

#### Spot cluster annotation

Each parental cluster’s mean cell type fraction value (theta values from BayesPrism output) for each cell type was calculated, and a template cell type fraction vector for each cluster was created. For each Visium spot in the untreated lymphomoid, Jennson-Shannon distance between the deconvolved cell type fraction vector and each of the 5 parental cluster templates were calculated. The spot was then attributed to that parental cluster whose template had the least distance from the spot’s cell type fraction.

#### Targeted mutation calling using exon sequencing

Genomic DNA was extracted from paraffin embedded tissue sections using the Generead DNA FFPE kit (Qiagen) following the manufacturer’s instructions. Whole exome sequencing was performed using Illumina NovaSeq 2zx150bp. Whole exome sequencing data for each sample was processed following the Genome Analysis Toolkit (GATK, v4.0.10.1) workflows and best practices^66^. Quality check was performed on each FASTQ file using FastQC^67^. Paired-end reads were then aligned the reference human genome (GRCh38) using bwa mem^68^ (v0.7.17, parameters: -M -v). SAM files were converted to BAM, sorted and indexed using samtools^69^ v0.1.19. Duplicate removal and read grouping were performed with Picard tools, and BAM files were further processed with GATK ‘BaseRecalibrator’ and ‘ApplyBQSR’. Mutation calling was performed using GATK ‘Mutect2’ in tumor-only mode, with the following parameters: ‘-L default_exome_hg38_refGene.bed --germline-resource af-only-gnomad.hg38.vcf.gz --af-of-alleles-not-in-resource 0.0000025 --disable-read-filter MateOnSameContigOrNoMappedMateReadFilter --panel-of-normals somatic-hg38_1000g_pon.hg38.vcf.gz’ (resources were retrieved from GATK bundle). Mutation filtering was performed with GATK ‘FilterMutectCalls’ and ‘FilterByOrientationBias’. The resulting VCF files were converted into MAF with vcf2maf v1.6.16. Oncogenic variants in the MAF files were then annotated using OncoKB^70^ ‘MafAnnotator’ tool. Variants were retained if they passed Mutect2 filtering or if they were annotated as oncogenic, and the latter were individually inspected.

### QUANTIFICATION AND STATISTICAL ANALYSIS

#### Similarity Index

We represented the intra-tumor cell type heterogeneity of each tumor sample or control lymphomoid *i* with a compositional vector 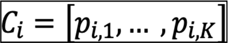, with 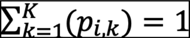, whose elements are the cell type proportions of the *K* cell types (non-proliferating CD20+ B cells, proliferating CD20+ B cells, CD4+ T cells, CD8+ T cells, CD68+ macrophages, otherCells) detected from the immunofluorescence image analyses. Then, we computed a Similarity Index between the compositional vectors of each pair of samples *i,j* as

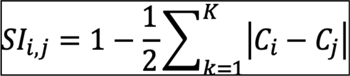

Finally, we computed the average SI of samples from the same patient (‘intra’, one data point per patient and per sample type i.e., tumor or control lymphomoid) and the average SI of samples belonging to different patients (‘inter’, one data point for each combination of patients x sample types). We tested the differences between ‘intra’ and ‘inter’ SI distributions computed for lymphomoids vs lymphomoids, lymphomoids vs tumors, and tumors vs tumors with Wilcoxon Rank-Sum test.

#### Statistics

Data were analyzed using R^71^ and the GRAPHPAD PRISM software. Normality was assessed using D’Agostino-Pearson’s omnibus normality test. Means were compared with two-tailed unpaired Student’s t-test or Mann-Whitney’s non-parametric test. One-way analysis of variance was performed to compare more than two variables. In case groups would not pass normality or data was comprised between 0 and 1, samples were analyzed using the Kruskal– Wallis test. To isolate differences between groups, the Dunn’s multiple comparisons test was performed. P-values are indicated for each experiment. Correlation coefficients were computed on weighted ratios dividing the sum of the values for each parameter (CD20, CD4, CD8 and CD68) by the sum of the total of cell counts, and the sum of CD20+KI67+ by the sum of CD20 counts. Likewise, the summaries presented in Figure 3c and d were computed on the same weighted ratios comparing the values of the treated lymphomoids to the untreated ones for each patient. Error bars indicate standard deviation unless stated otherwise.

## Acknowledgments

We thank all the patients that have participated in this study. We also thank the histology, bioimaging, and optics platform, the flow cytometry, and the gene expression facilities at EPFL for their technical support, Dr. Olaia Naveiras for the infrastructure, and Dr. Gonzalez and Dr. Perentes for their participation in the collection of surgical biopsies. This work has been supported by Fondation Aclon, Accentus Foundation, and the Fondazione San Salvatore.

## Author contributions

A.S.M. methods development, analysis, J.E. acquisition of clinical samples, D.S., D.T., M.V. computational analyses, D.M., I. L. pathology assessment, T.K., R.D. and A.C. accessibility of human samples, G.C. supervision of the computational analyses, A.S.M. and E.O. designed and supervised the study, E.O. A.S.M. J.E and D.S. wrote the manuscript with input from all the authors.

## Supplemental information

Document Supplemental_information, Table S1, Table S2, Table S3, Table S4, Table S5.

## References

1. Black, J.R.M., and McGranahan, N. (2021). Genetic and non-genetic clonal diversity in cancer evolution. Nat Rev Cancer 21, 379–392. 10.1038/s41568-021-00336-2.

2. Barroso, M., Chheda, M.G., Clevers, H., Elez, E., Kaochar, S., Kopetz, S.E., Li, X.-N., Meric-Bernstam, F., Meyer, C.A., Mou, H., et al. (2022). A path to translation: How 3D patient tumor avatars enable next generation precision oncology. Cancer Cell, S1535610822004755. 10.1016/j.ccell.2022.09.017.

3. Abdolahi, S., Ghazvinian, Z., Muhammadnejad, S., Saleh, M., Asadzadeh Aghdaei, H., and Baghaei, K. (2022). Patient-derived xenograft (PDX) models, applications and challenges in cancer research. J Transl Med 20, 206. 10.1186/s12967-022-03405-8.

4. LeBlanc, V.G., Trinh, D.L., Aslanpour, S., Hughes, M., Livingstone, D., Jin, D., Ahn, B.Y., Blough, M.D., Cairncross, J.G., Chan, J.A., et al. (2022). Single-cell landscapes of primary glioblastomas and matched explants and cell lines show variable retention of inter- and intratumor heterogeneity. Cancer Cell 40, 379–392.e9. 10.1016/j.ccell.2022.02.016.

5. Voabil, P., de Bruijn, M., Roelofsen, L.M., Hendriks, S.H., Brokamp, S., van den Braber, M., Broeks, A., Sanders, J., Herzig, P., Zippelius, A., et al. (2021). An ex vivo tumor fragment platform to dissect response to PD-1 blockade in cancer. Nat Med 27, 1250–1261. 10.1038/s41591-021-01398-3.

6. Broutier, L., Mastrogiovanni, G., Verstegen, M.M., Francies, H.E., Gavarró, L.M., Bradshaw, C.R., Allen, G.E., Arnes-Benito, R., Sidorova, O., Gaspersz, M.P., et al. (2017). Human primary liver cancer-derived organoid cultures for disease modeling and drug screening. Nat Med 23, 1424–1435. 10.1038/nm.4438.

7. Ganesh, K., Wu, C., O’Rourke, K.P., Szeglin, B.C., Zheng, Y., Sauvé, C.-E.G., Adileh, M., Wasserman, I., Marco, M.R., Kim, A.S., et al. (2019). A rectal cancer organoid platform to study individual responses to chemoradiation. Nat Med 25, 1607–1614. 10.1038/s41591-019-0584-2.

8. Tiriac, H., Belleau, P., Engle, D.D., Plenker, D., Deschênes, A., Somerville, T.D.D., Froeling, F.E.M., Burkhart, R.A., Denroche, R.E., Jang, G.-H., et al. (2018). Organoid Profiling Identifies Common Responders to Chemotherapy in Pancreatic Cancer. Cancer Discov 8, 1112–1129. 10.1158/2159-8290.CD-18-0349.

9. Yao, Y., Xu, X., Yang, L., Zhu, J., Wan, J., Shen, L., Xia, F., Fu, G., Deng, Y., Pan, M., et al. (2020). Patient-Derived Organoids Predict Chemoradiation Responses of Locally Advanced Rectal Cancer. Cell Stem Cell 26, 17–26.e6. 10.1016/j.stem.2019.10.010.

10. Khan, A.O., Rodriguez-Romera, A., Reyat, J.S., Olijnik, A.-A., Colombo, M., Wang, G., Wen, W.X., Sousos, N., Murphy, L.C., Grygielska, B., et al. (2022). Human bone marrow organoids for disease modelling, discovery and validation of therapeutic targets in hematological malignancies. Cancer Discovery, CD-22-0199. 10.1158/2159-8290.CD-22-0199.

11. Guillen, K.P., Fujita, M., Butterfield, A.J., Scherer, S.D., Bailey, M.H., Chu, Z., DeRose, Y.S., Zhao, L., Cortes-Sanchez, E., Yang, C.-H., et al. (2022). A human breast cancer-derived xenograft and organoid platform for drug discovery and precision oncology. Nat Cancer 3, 232–250. 10.1038/s43018-022-00337-6.

12. Decaup, E., Jean, C., Laurent, C., Gravelle, P., Fruchon, S., Capilla, F., Marrot, A., Al Saati, T., Frenois, F.-X., Laurent, G., et al. (2013). Anti-tumor activity of obinutuzumab and rituximab in a follicular lymphoma 3D model. Blood Cancer J 3, e131. 10.1038/bcj.2013.32.

13. Mannino, R.G., Santiago-Miranda, A.N., Pradhan, P., Qiu, Y., Mejias, J.C., Neelapu, S.S., Roy, K., and Lam, W.A. (2017). 3D microvascular model recapitulates the diffuse large B-cell lymphoma tumor microenvironment in vitro. Lab Chip 17, 407–414. 10.1039/c6lc01204c.

14. Tian, Y.F., Ahn, H., Schneider, R.S., Yang, S.N., Roman-Gonzalez, L., Melnick, A.M., Cerchietti, L., and Singh, A. (2015). Integrin-specific hydrogels as adaptable tumor organoids for malignant B and T cells. Biomaterials 73, 110–119. 10.1016/j.biomaterials.2015.09.007.

15. Foxall, R., Narang, P., Glaysher, B., Hub, E., Teal, E., Coles, M.C., Ashton-Key, M., Beers, S.A., and Cragg, M.S. (2021). Developing a 3D B Cell Lymphoma Culture System to Model Antibody Therapy. Front. Immunol. 11, 605231. 10.3389/fimmu.2020.605231.

16. Faria, C., Gava, F., Gravelle, P., Valero, J.G., Dobaño-López, C., Acker, N.V., Quelen, C., Jalowicki, G., Morin, R., Rossi, C., et al. (2023). Patient-derived lymphoma spheroids integrating immune tumor microenvironment as preclinical follicular lymphoma models for personalized medicine. J Immunother Cancer 11, e007156. 10.1136/jitc-2023-007156.

17. Shah, S.B., Carlson, C.R., Lai, K., Zhong, Z., Marsico, G., Lee, K.M., Félix Vélez, N.E., Abeles, E.B., Allam, M., Hu, T., et al. (2023). Combinatorial treatment rescues tumour-microenvironment-mediated attenuation of MALT1 inhibitors in B-cell lymphomas. Nat. Mater. 22, 511–523. 10.1038/s41563-023-01495-3.

18. Araujo-Ayala, F., Dobaño-López, C., Valero, J.G., Nadeu, F., Gava, F., Faria, C., Norlund, M., Morin, R., Bernes-Lasserre, P., Serrat, N., et al. (2023). A novel patient-derived 3D model recapitulates mantle cell lymphoma lymph node signaling, immune profile and in vivo ibrutinib responses. Leukemia 37, 1311–1323. 10.1038/s41375-023-01885-1.

19. Kastenschmidt, J.M., Schroers-Martin, J.G., Sworder, B.J., Sureshchandra, S., Khodadoust, M.S., Liu, C.L., Olsen, M., Kurtz, D.M., Diehn, M., Wagar, L.E., et al. (2024). A human lymphoma organoid model for evaluating and targeting the follicular lymphoma tumor immune microenvironment. Cell Stem Cell 31, 410–420.e4. 10.1016/j.stem.2024.01.012.

20. Pasqualucci, L., and Dalla-Favera, R. (2018). Genetics of diffuse large B-cell lymphoma. Blood 131, 2307–2319. 10.1182/blood-2017-11-764332.

21. Dheilly, E., Battistello, E., Katanayeva, N., Sungalee, S., Michaux, J., Duns, G., Wehrle, S., Sordet-Dessimoz, J., Mina, M., Racle, J., et al. (2020). Cathepsin S Regulates Antigen Processing and T Cell Activity in Non-Hodgkin Lymphoma. Cancer Cell 37, 674–689.e12. 10.1016/j.ccell.2020.03.016.

22. Morin, R.D., Mendez-Lago, M., Mungall, A.J., Goya, R., Mungall, K.L., Corbett, R.D., Johnson, N.A., Severson, T.M., Chiu, R., Field, M., et al. (2011). Frequent mutation of histone-modifying genes in non-Hodgkin lymphoma. Nature 476, 298–303. 10.1038/nature10351.

23. Pasqualucci, L., Dominguez-Sola, D., Chiarenza, A., Fabbri, G., Grunn, A., Trifonov, V., Kasper, L.H., Lerach, S., Tang, H., Ma, J., et al. (2011). Inactivating mutations of acetyltransferase genes in B-cell lymphoma. Nature 471, 189–195. 10.1038/nature09730.

24. Pasqualucci, L., Trifonov, V., Fabbri, G., Ma, J., Rossi, D., Chiarenza, A., Wells, V.A., Grunn, A., Messina, M., Elliot, O., et al. (2011). Analysis of the coding genome of diffuse large B-cell lymphoma. Nat Genet 43, 830–837. 10.1038/ng.892.

25. Chambwe, N., Kormaksson, M., Geng, H., De, S., Michor, F., Johnson, N.A., Morin, R.D., Scott, D.W., Godley, L.A., Gascoyne, R.D., et al. (2014). Variability in DNA methylation defines novel epigenetic subgroups of DLBCL associated with different clinical outcomes. Blood 123, 1699–1708. 10.1182/blood-2013-07-509885.

26. Zhang, J., Vlasevska, S., Wells, V.A., Nataraj, S., Holmes, A.B., Duval, R., Meyer, S.N., Mo, T., Basso, K., Brindle, P.K., et al. (2017). The CREBBP Acetyltransferase Is a Haploinsufficient Tumor Suppressor in B-cell Lymphoma. Cancer Discovery 7, 322–337. 10.1158/2159-8290.CD-16-1417.

27. Jiang, Y., Ortega-Molina, A., Geng, H., Ying, H.-Y., Hatzi, K., Parsa, S., McNally, D., Wang, L., Doane, A.S., Agirre, X., et al. (2017). *CREBBP* Inactivation Promotes the Development of HDAC3-Dependent Lymphomas. Cancer Discovery 7, 38–53. 10.1158/2159-8290.CD-16-0975.

28. Alizadeh, A.A., Eisen, M.B., Davis, R.E., Ma, C., Lossos, I.S., Rosenwald, A., Boldrick, J.C., Sabet, H., Tran, T., Yu, X., et al. (2000). Distinct types of diffuse large B-cell lymphoma identified by gene expression profiling. Nature 403, 503–511. 10.1038/35000501.

29. Lu, X., Fernando, T.M., Lossos, C., Yusufova, N., Liu, F., Fontán, L., Durant, M., Geng, H., Melnick, J., Luo, Y., et al. (2018). PRMT5 interacts with the BCL6 oncoprotein and is required for germinal center formation and lymphoma cell survival. Blood 132, 2026–2039. 10.1182/blood-2018-02-831438.

30. Carbone, A., Roulland, S., Gloghini, A., Younes, A., von Keudell, G., López-Guillermo, A., and Fitzgibbon, J. (2019). Follicular lymphoma. Nat Rev Dis Primers 5, 83. 10.1038/s41572-019-0132-x.

31. Sehn, L.H., and Salles, G. (2021). Diffuse Large B-Cell Lymphoma. N Engl J Med 384, 842–858. 10.1056/NEJMra2027612.

32. de Leval, L., and Jaffe, E.S. (2020). Lymphoma Classification. Cancer J 26, 176–185. 10.1097/PPO.0000000000000451.

33. Alaggio, R., Amador, C., Anagnostopoulos, I., Attygalle, A.D., Araujo, I.B. de O., Berti, E., Bhagat, G., Borges, A.M., Boyer, D., Calaminici, M., et al. (2022). The 5th edition of the World Health Organization Classification of Haematolymphoid Tumours: Lymphoid Neoplasms. Leukemia, 1–29. 10.1038/s41375-022-01620-2.

34. Campo, E., Jaffe, E.S., Cook, J.R., Quintanilla-Martinez, L., Swerdlow, S.H., Anderson, K.C., Brousset, P., Cerroni, L., De Leval, L., Dirnhofer, S., et al. (2022). The International Consensus Classification of Mature Lymphoid Neoplasms: a report from the Clinical Advisory Committee. Blood 140, 1229–1253. 10.1182/blood.2022015851.

35. Chapuy, B., Stewart, C., Dunford, A.J., Kim, J., Kamburov, A., Redd, R.A., Lawrence, M.S., Roemer, M.G.M., Li, A.J., Ziepert, M., et al. (2018). Molecular subtypes of diffuse large B cell lymphoma are associated with distinct pathogenic mechanisms and outcomes. Nat Med 24, 679–690. 10.1038/s41591-018-0016-8.

36. Schmitz, R., Wright, G.W., Huang, D.W., Johnson, C.A., Phelan, J.D., Wang, J.Q., Roulland, S., Kasbekar, M., Young, R.M., Shaffer, A.L., et al. (2018). Genetics and Pathogenesis of Diffuse Large B-Cell Lymphoma. New England Journal of Medicine 378, 1396–1407. 10.1056/NEJMoa1801445.

37. Basso, K., and Dalla-Favera, R. (2015). Germinal centres and B cell lymphomagenesis. Nat Rev Immunol 15, 172–184. 10.1038/nri3814.

38. Scott, D.W., and Gascoyne, R.D. (2014). The tumour microenvironment in B cell lymphomas. Nat Rev Cancer 14, 517–534. 10.1038/nrc3774.

39. Pangault, C., Amé-Thomas, P., Ruminy, P., Rossille, D., Caron, G., Baia, M., De Vos, J., Roussel, M., Monvoisin, C., Lamy, T., et al. (2010). Follicular lymphoma cell niche: identification of a preeminent IL-4-dependent T(FH)-B cell axis. Leukemia 24, 2080–2089. 10.1038/leu.2010.223.

40. Amé-Thomas, P., and Tarte, K. (2014). The yin and the yang of follicular lymphoma cell niches: role of microenvironment heterogeneity and plasticity. Semin Cancer Biol 24, 23–32. 10.1016/j.semcancer.2013.08.001.

41. PDQ Adult Treatment Editorial Board (2002). Adult Non-Hodgkin Lymphoma Treatment (PDQ®): Health Professional Version. In PDQ Cancer Information Summaries (National Cancer Institute (US)).

42. Sheikh, S., Migliorini, D., and Lang, N. (2022). CAR T-Based Therapies in Lymphoma: A Review of Current Practice and Perspectives. Biomedicines 10, 1960. 10.3390/biomedicines10081960.

43. Mussetti, A., and Sureda, A. (2022). Second-line CAR T cells for lymphomas. The Lancet 399, 2247–2249. 10.1016/S0140-6736(22)00790-5.

44. Chaudhari, K., Rizvi, S., and Syed, B.A. (2019). Non-Hodgkin lymphoma therapy landscape. Nat Rev Drug Discov 18, 663–664. 10.1038/d41573-019-00051-6.

45. Younes, A., Ansell, S., Fowler, N., Wilson, W., de Vos, S., Seymour, J., Advani, R., Forero, A., Morschhauser, F., Kersten, M.J., et al. (2017). The landscape of new drugs in lymphoma. Nat Rev Clin Oncol 14, 335–346. 10.1038/nrclinonc.2016.205.

46. Neal, J.T., Li, X., Zhu, J., Giangarra, V., Grzeskowiak, C.L., Ju, J., Liu, I.H., Chiou, S.-H., Salahudeen, A.A., Smith, A.R., et al. (2018). Organoid Modeling of the Tumor Immune Microenvironment. Cell 175, 1972–1988.e16. 10.1016/j.cell.2018.11.021.

47. Egle, A., Harris, A.W., Bath, M.L., O’Reilly, L., and Cory, S. (2004). VavP-Bcl2 transgenic mice develop follicular lymphoma preceded by germinal center hyperplasia. Blood 103, 2276–2283. 10.1182/blood-2003-07-2469.

48. Zhang, X., Lan, Y., Xu, J., Quan, F., Zhao, E., Deng, C., Luo, T., Xu, L., Liao, G., Yan, M., et al. (2019). CellMarker: a manually curated resource of cell markers in human and mouse. Nucleic Acids Research 47, D721–D728. 10.1093/nar/gky900.

49. Reimold, A.M., Iwakoshi, N.N., Manis, J., Vallabhajosyula, P., Szomolanyi-Tsuda, E., Gravallese, E.M., Friend, D., Grusby, M.J., Alt, F., and Glimcher, L.H. (2001). Plasma cell differentiation requires the transcription factor XBP-1. Nature 412, 300–307. 10.1038/35085509.

50. Shi, W., Liao, Y., Willis, S.N., Taubenheim, N., Inouye, M., Tarlinton, D.M., Smyth, G.K., Hodgkin, P.D., Nutt, S.L., and Corcoran, L.M. (2015). Transcriptional profiling of mouse B cell terminal differentiation defines a signature for antibody-secreting plasma cells. Nat Immunol 16, 663–673. 10.1038/ni.3154.

51. Nutt, S.L., and Kee, B.L. (2007). The Transcriptional Regulation of B Cell Lineage Commitment. Immunity 26, 715–725. 10.1016/j.immuni.2007.05.010.

52. Hwang, I.-Y., Park, C., Harrison, K., and Kehrl, J.H. (2019). Biased S1PR1 Signaling in B Cells Subverts Responses to Homeostatic Chemokines, Severely Disorganizing Lymphoid Organ Architecture. The Journal of Immunology 203, 2401–2414. 10.4049/jimmunol.1900678.

53. Hodson, D.J., Shaffer, A.L., Xiao, W., Wright, G.W., Schmitz, R., Phelan, J.D., Yang, Y., Webster, D.E., Rui, L., Kohlhammer, H., et al. (2016). Regulation of normal B-cell differentiation and malignant B-cell survival by OCT2. Proc. Natl. Acad. Sci. U.S.A. 113. 10.1073/pnas.1600557113.

54. Chu, T., Wang, Z., Pe’er, D., and Danko, C.G. (2022). Cell type and gene expression deconvolution with BayesPrism enables Bayesian integrative analysis across bulk and single-cell RNA sequencing in oncology. Nat Cancer 3, 505–517. 10.1038/s43018-022-00356-3.

55. Kleshchevnikov, V., Shmatko, A., Dann, E., Aivazidis, A., King, H.W., Li, T., Elmentaite, R., Lomakin, A., Kedlian, V., Gayoso, A., et al. (2022). Cell2location maps fine-grained cell types in spatial transcriptomics. Nat Biotechnol 40, 661–671. 10.1038/s41587-021-01139-4.

56. Battistello, E., Katanayeva, N., Dheilly, E., Tavernari, D., Donaldson, M.C., Bonsignore, L., Thome, M., Christie, A.L., Murakami, M.A., Michielin, O., et al. (2018). Pan-SRC kinase inhibition blocks B-cell receptor oncogenic signaling in non-Hodgkin lymphoma. Blood 131, 2345–2356. 10.1182/blood-2017-10-809210.

57. Noy, A., de Vos, S., Coleman, M., Martin, P., Flowers, C.R., Thieblemont, C., Morschhauser, F., Collins, G.P., Ma, S., Peles, S., et al. (2020). Durable ibrutinib responses in relapsed/refractory marginal zone lymphoma: long-term follow-up and biomarker analysis. Blood Advances 4, 5773–5784. 10.1182/bloodadvances.2020003121.

58. Laursen, M.B., Falgreen, S., Bødker, J.S., Schmitz, A., Kjeldsen, M.K., Sørensen, S., Madsen, J., El-Galaly, T.C., Bøgsted, M., Dybkær, K., et al. (2014). Human B-cell cancer cell lines as a preclinical model for studies of drug effect in diffuse large B-cell lymphoma and multiple myeloma. Experimental Hematology 42, 927–938. 10.1016/j.exphem.2014.07.263.

59. Chapuy, B., Cheng, H., Watahiki, A., Ducar, M.D., Tan, Y., Chen, L., Roemer, M.G.M., Ouyang, J., Christie, A.L., Zhang, L., et al. (2016). Diffuse large B-cell lymphoma patient-derived xenograft models capture the molecular and biological heterogeneity of the disease. Blood 127, 2203–2213. 10.1182/blood-2015-09-672352.

60. Zhang, L., Nomie, K., Zhang, H., Bell, T., Pham, L., Kadri, S., Segal, J., Li, S., Zhou, S., Santos, D., et al. (2017). B-Cell Lymphoma Patient-Derived Xenograft Models Enable Drug Discovery and Are a Platform for Personalized Therapy. Clin Cancer Res 23, 4212–4223. 10.1158/1078-0432.CCR-16-2703.

61. Hilton, L.K., Ngu, H.S., Collinge, B., Dreval, K., Ben-Neriah, S., Rushton, C.K., Wong, J.C.H., Cruz, M., Roth, A., Boyle, M., et al. (2023). Relapse Timing Is Associated With Distinct Evolutionary Dynamics in Diffuse Large B-Cell Lymphoma. J Clin Oncol 41, 4164–4177. 10.1200/JCO.23.00570.

62. Wagar, L.E., Salahudeen, A., Constantz, C.M., Wendel, B.S., Lyons, M.M., Mallajosyula, V., Jatt, L.P., Adamska, J.Z., Blum, L.K., Gupta, N., et al. (2021). Modeling human adaptive immune responses with tonsil organoids. Nat Med 27, 125–135. 10.1038/s41591-020-01145-0.

63. Bankhead, P., Loughrey, M.B., Fernández, J.A., Dombrowski, Y., McArt, D.G., Dunne, P.D., McQuaid, S., Gray, R.T., Murray, L.J., Coleman, H.G., et al. (2017). QuPath: Open source software for digital pathology image analysis. Sci Rep 7, 16878. 10.1038/s41598-017-17204-5.

64. Greenwald, N.F., Miller, G., Moen, E., Kong, A., Kagel, A., Dougherty, T., Fullaway, C.C., McIntosh, B.J., Leow, K.X., Schwartz, M.S., et al. (2022). Whole-cell segmentation of tissue images with human-level performance using large-scale data annotation and deep learning. Nat Biotechnol 40, 555–565. 10.1038/s41587-021-01094-0.

65. Schapiro, D., Sokolov, A., Yapp, C., Chen, Y.-A., Muhlich, J.L., Hess, J., Creason, A.L., Nirmal, A.J., Baker, G.J., Nariya, M.K., et al. (2022). MCMICRO: a scalable, modular image-processing pipeline for multiplexed tissue imaging. Nat Methods 19, 311–315. 10.1038/s41592-021-01308-y.

66. Auwera, G.A., Carneiro, M.O., Hartl, C., Poplin, R., del Angel, G., Levy-Moonshine, A., Jordan, T., Shakir, K., Roazen, D., Thibault, J., et al. (2013). From FastQ Data to High-Confidence Variant Calls: The Genome Analysis Toolkit Best Practices Pipeline. Current Protocols in Bioinformatics 43. 10.1002/0471250953.bi1110s43.

67. Babraham Bioinformatics - FastQC A Quality Control tool for High Throughput Sequence Data https://www.bioinformatics.babraham.ac.uk/projects/fastqc/.

68. Li, H. (2013). Aligning sequence reads, clone sequences and assembly contigs with BWA-MEM. 10.48550/ARXIV.1303.3997.

69. Twelve years of SAMtools and BCFtools | GigaScience | Oxford Academic https://academic.oup.com/gigascience/article/10/2/giab008/6137722?login=false.

70. Chakravarty, D., Gao, J., Phillips, S., Kundra, R., Zhang, H., Wang, J., Rudolph, J.E., Yaeger, R., Soumerai, T., Nissan, M.H., et al. (2017). OncoKB: A Precision Oncology Knowledge Base. JCO Precision Oncology, 1–16. 10.1200/PO.17.00011.

71. R Core Team (2023) R: A Language and Environment for Statistical Computing.

